# Impact of spatial organization on a novel auxotrophic interaction among soil microbes

**DOI:** 10.1101/195339

**Authors:** Xue Jiang, Christian Zerfaß, Ruth Eichmann, Munehiro Asally, Patrick Schäfer, Orkun S Soyer

## Abstract

A key prerequisite to achieve a deeper understanding of microbial communities and to engineer synthetic ones is to identify the individual metabolic interactions among key species and how these interactions are affected by different environmental factors. Deciphering the physiological basis of species-species and species-environment interactions in spatially organized environment requires reductionist approaches using ecologically and functionally relevant species. To this end, we focus here on a specific defined system to study the metabolic interactions in a spatial context among a plant-beneficial endophytic fungus *Serendipita indica,* and the soil-dwelling model bacterium *Bacillus subtilis*. Focusing on the growth dynamics of *S. indica* under defined conditions, we identified an auxotrophy in this organism for thiamine, which is a key co-factor for essential reactions in the central carbon metabolism. We found that *S. indica* growth is restored in thiamine-free media, when co-cultured with *B. subtilis*. The success of this auxotrophic interaction, however, was dependent on the spatial and temporal organization of the system; the beneficial impact of *B. subtilis* was only visible when its inoculation was separated from that of *S. indica* either in time or space. These findings describe a key auxotrophic interaction in the soil among organisms that are shown to be important for plant ecosystem functioning, and point to the potential importance of spatial and temporal organization for the success of auxotrophic interactions. These points can be particularly important for engineering of minimal functional synthetic communities as plant-seed treatments and for vertical farming under defined conditions.

## Introduction

Higher-level functions and population dynamics within microbial communities are underpinned by the interactions among the composing species within the community and their environment (Falkowski *et al.*, 2008; Sañudo-Wilhelmy *et al.*, 2014). Deciphering these interactions is a pre-requisite to understand and manage complex natural communities (Abreu and Taga, 2016; Widder *et al.*, 2016) and to achieve community-level synthetic engineering (Großkopf and Soyer, 2014; Hays *et al.*, 2015; Lindemann *et al.*, 2016). To this end, increasing numbers of experimental studies and (meta)genomic surveys have shown that auxotrophic interactions, involving vitamins and amino acids, are wide-spread in many microbial natural communities (Sañudo-Wilhelmy *et al.*, 2014; Morris *et al.*, 2012; Helliwell *et al.*, 2013) and can also be engineered genetically to create synthetic communities (Mee *et al.*, 2014; Pande *et al.*, 2014). Specific auxotrophic interactions among microbes are shown to influence ecosystem functioning; e.g. infection outcomes within higher organisms (Wargo and Hogan, 2006), ecological population dynamics in the oceans (Sañudo-Wilhelmy *et al.*, 2014), and the level of biodegradation of organic matter under anoxic conditions (Schink, 1997; Embree *et al.*, 2015).

It has been suggested that auxotrophies can result from reduced selective pressures for maintaining biosynthesis capabilities under stable metabolite availability due to abiotic or biotic sources (Morris *et al.*, 2012; Helliwell *et al.*, 2013). This proposition is supported by the observed independent evolution of vitamin and amino acid auxotrophies in different, unrelated taxa (Helliwell *et al.*, 2011; Rodionova *et al.*, 2015), and points to a direct linkage between ecological dynamics and evolution of auxotrophies (Embree *et al.*, 2015). The possible fluctuations in a metabolite’s availability in time and space would be expected to impact both the emergence of auxotrophies and the population dynamics of resulting auxotrophic species. For example, in the marine environment, where the observed auxotrophies relate mostly to the loss of biosynthesis capacity for vitamins and amino acids, population dynamics of auxotrophic species are believed to be directly linked to those of ‘provider’ species (Helliwell *et al.*, 2013; Hom and Murray, 2014; Sañudo-Wilhelmy *et al.*, 2014). The ecological influences of auxotrophic species on community structure and population dynamics can also be exerted by abiotic fluctuations or directly by the abundances and actions of higher organisms within the system.

These ecological influences on microbial population dynamics can increase significantly in spatially organized systems. Yet, the spatial context of microbial interactions is under-studied. Considering that each species can display multiple metabolic actions that all affect a common environment, it is not clear if auxotrophic interactions can always be successfully established even if genetic/metabolic complementarity is present. For example, changes in environmental pH upon growth of one species can affect its subsequent interactions with other microbes (Vylkova, 2017; Ratzke and Gore, 2017). Similarly, extensive oxygen depletion upon microbial colony growth plates (Peters and Wimpenny, 1987; Dietrich *et al.*, 2013; Kempes and Okegbe, 2014) can directly influence subsequent or simultaneous growth of different species or their interactions. These possible interplays between species-species and species-environment interactions are currently not well-understood and only explored in few studies, which used either synthetically engineered interactions (Shou *et al.*, 2007; Mee *et al.*, 2014) or enriched microbial communities (Embree *et al.*, 2015). This lack of understanding, however, causes a particular challenge for the engineering of novel applications of microbial communities with inherent spatial organisation, such as seen in agriculture and involving for example closed-ecosystem production, seed treatment and microbe-based biofertilisation (Gòdia *et al.*, 2002; Lucy *et al.*, 2004; Richardson *et al.*, 2011).

Towards addressing this challenge, we focus here on identifying potential metabolic interactions among plant-beneficial endophytic fungus *Serendipita indica* (previously called *Piriformospora indica* (Weiss *et al.*, 2016)) and the common soil microbe *Bacillus subtilis.* Identifying a defined media for *S. indica*, we found it to be auxotrophic for the vitamin B1, thiamine. To study the potential auxotrophic interactions of *S. indica*, we then created a co-culture system using *Bacillus subtilis*. We found that *S. indica* thiamine auxotrophy can be satisfied and its growth is restored in the presence of *B. subtilis*. The success of this auxotrophic interaction, however, is strongly dependent on temporal and spatial organization in the system. These findings and the established synthetic co-culture can act as a basis to develop a more complete functional synthetic community, as advocated for biotechnological applications and for gaining insights into community function (Mee and Wang, 2012; Großkopf and Soyer, 2014; Widder *et al.*, 2016; Lindemann *et al.*, 2016). With the inclusion of a plant, such a synthetic community can allow further insights into microbe-microbe, and microbe-plant interactions and development of new agricultural technologies such as in seed coating and vertical farming in controlled environments.

## Results

### *S. indica* is auxotrophic for thiamine

*S. indica* is an endophytic fungus that can colonize roots of a wide range of plants and can confer a range of beneficial effects, including enhancing plant growth, resistance to biotic and abiotic stresses (Waller *et al.*, 2005; Sherameti *et al.*, 2008; Vadassery *et al.*, 2009), promotion of adventitious root formation in cuttings (Druege *et al.*, 2007), and assisting phosphate assimilation (Yadav *et al.*, 2010). Despite its broad host range, *S. indica* also has the ability to grow in the absence of host plants (Kumar *et al.*, 2011). Exploiting this ability, we attempted to create a fully defined growth medium that was based on previous physiological studies on *S. indica* (Zuccaro *et al.*, 2011; Kumar *et al.*, 2011; Jacobs *et al.*, 2011; Varma *et al.*, 2012; Qiang *et al.*, 2012). Using our defined media, we tested the effect of the vitamins on growth by cultivating *S. indica* in a series of vitamin-free media each supplemented by a specific vitamin (Figure 1). The analyses showed that *S. indica* is auxotrophic for thiamine; while none of the other individual vitamin additions supported growth, thiamine and full vitamin addition did. This finding was further confirmed by growing *S. indica* on plates supplemented with an additional agar block only containing a defined amount of thiamine. In this case, growth of *S. indica* resulted in expansion towards the thiamine agar block, suggesting that growth occurs on a thiamine gradient or is linked with an active chemotaxis towards thiamine source (Figure S1). We also quantified the growth of *S. indica* with different concentrations of thiamine and found that hyphae growth and spore formation showed a positive, but saturating, dependency on thiamine levels (Figure S2). At zero concentration of thiamine in the media, we still observe germination and very little hyphal growth (Figure S2), possibly supported by thiamine stored in spores. Besides these measurements, we also observed thiamine effect on *S. indica* growth using time-lapse microscopy (see further discussion below and supplementary videos 1 and 2).

**Figure 1.**
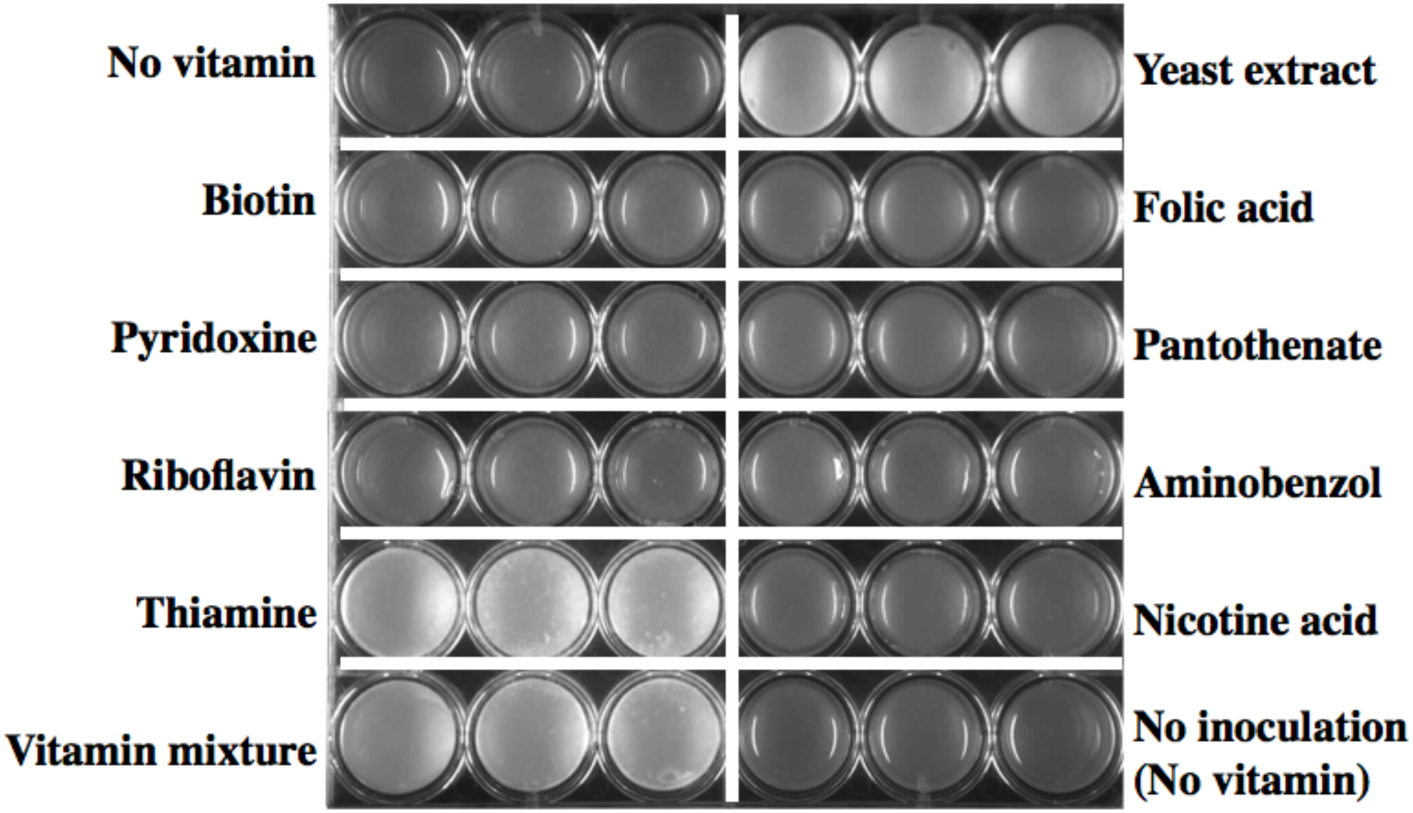
*S. indica* growth under different conditions. Growth on agar plates with defined medium supplemented with different vitamins as shown on each row and column. Each treatment has three replicates presented in 3 adjacent wells. *S. indica* grows in white colonies. Images were taken after two weeks of growth.

### *S. indica* auxotrophy is reflected in its genomic enzyme content

To support and better understand these physiological results, we analyzed the *S. indica* genome for the presence of genes associated with thiamine use, biosynthesis, and transport. Bioinformatics analysis revealed that *S. indica* lacks most of the genes of the thiamine-biosynthesis pathway (Table S1, Figure 2a). In particular, we were not able to find any homologs of the genes *thi5* and *thi6*. The former encodes the enzyme involved in the synthesis of the thiamine-precursor hydroxymethylpyrimidine (HMP), while the later encodes the bifunctional enzyme acting as thiamine phosphate pyrophosphorylase and hydroxyethylthiazole kinase. A homolog of the gene *thi4*, the product of which also relates to mitochondrial DNA damage tolerance (Machado *et al.*, 1997; Hohmann and Meacock, 1998; Wightman, 2003), is present. We also found that *S. indica* contains a homolog of the *thi7* (or alternative name *thi10*) that encodes a thiamine transporter, and a homolog of the *pho3* gene, whose product catalyzes dephosphorylation of thiamine phosphate to thiamine, thereby increasing its uptake (Wightman, 2003). These findings suggest that *S. indica* is unable to synthesize thiamine, but can acquire thiamine from the environment, as also suggested for other microbes that encode thiamine salvage pathways (Jenkins *et al.*, 2007). The utilization of thiamine in physiology is evidenced by the presence of homologs of the *thi80* gene, which encodes a thiamine pyrophosphokinase involved in the catalysis of thiamine into thiamine pyrophosphate (ThPP), the presence of at least one gene encoding a ThPP binding domain-containing protein, and the role of ThPP as a key co-factor involved in central metabolic reactions (see Figure 2b). The last point involves key metabolic reactions such as pyruvate fermentation and conversion for entry into the citric acid cycle (TCA), oxo-glutarate to succinyl-CoA conversion in the TCA cycle, transketolase reactions in the pentose phosphate pathway, and biosynthesis reactions for leucine, isoleucine and valine (Michal and Schomburg, 1999)

**Figure 2.**
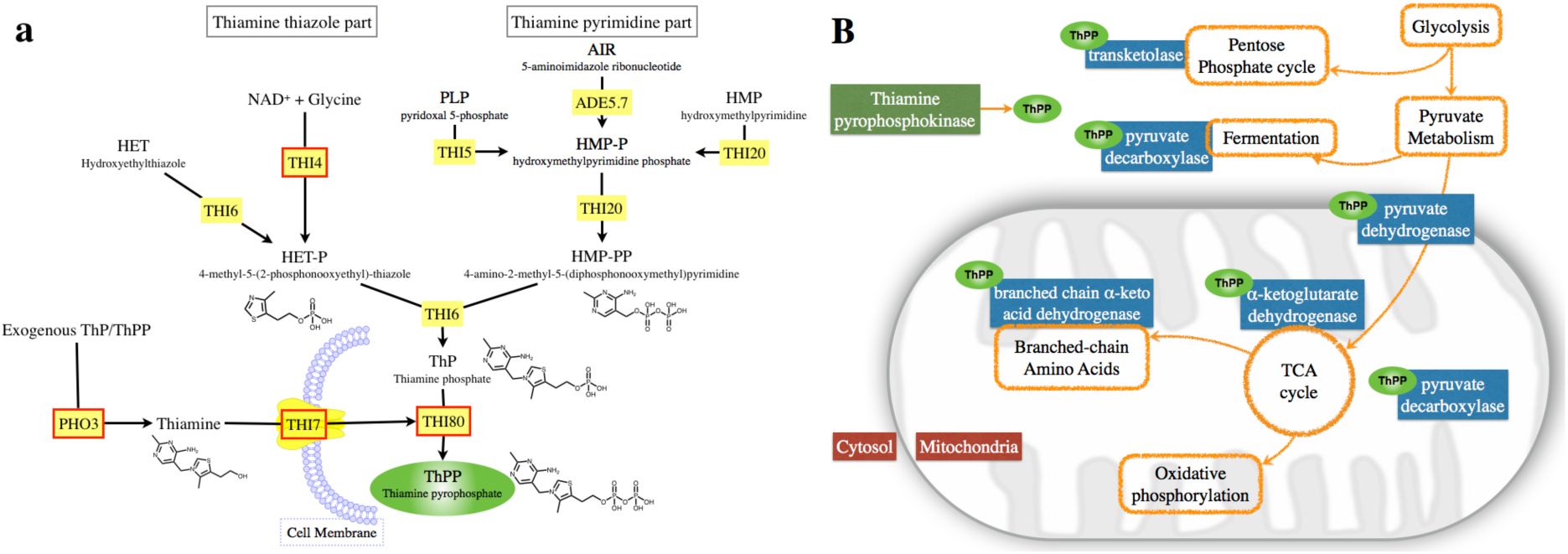
Thiamine related genes and reactions. **a.** Overview of the thiamine biosynthesis pathway in *Saccharomyces cerevisiae* based on (Wightman, 2003), and as included in the KEGG metabolic pathways (Pathway: sce00730) (Kanehisa and Goto, 2000). Yellow squares indicate genes encoding for the enzymes in the corresponding reactions. Red borders indicate genes for which there are *S. cerevisiae* homologues in *S. indica* (see *Methods*). **b.** Simplified schematic of central metabolism in eukaryotic cells with cytosol and mitochondria compartments indicated. Orange enclosures show reactions of the central metabolism, while blue squares show essential enzymes catalyzing these reactions and requiring ThPP as a co-factor (shown in green).

### *B. subtilis* complements *S. indica’s* auxotrophy for thiamine and promotes its growth

Given the crucial role of thiamine-derived co-factors in central metabolism, *S. indica* growth in nature apparently depends on environment-derived thiamine. Indeed, thiamine can be synthesized by various bacteria, fungi and plants (Begley *et al.*, 1999; Jenkins *et al.*, 2007; Jurgenson *et al.*, 2009). Among these, *B. subtilis*, a bacterium commonly found in soil (Hong *et al.*, 2009), is an established model organism (Mader *et al.*, 2011), and well-studied for thiamine biosynthesis (Schyns *et al.*, 2005; Begley *et al.*, 1999). Combined with the fact that *B. subtilis* is normally a plant-beneficial microbe (Castillo *et al.*, 2013), this motivated us to explore the possibility that the identified *S. indica* auxotrophy for thiamine could be satisfied upon co-culturing with *B. subtilis*. We created co-cultures of these two species on agar plates using our defined media and two common nitrogen sources to evaluate possible auxotrophic interaction under these conditions. We found that *B. subtilis* could indeed stimulate *S. indica* growth under thiamine-free conditions and that *S. indica* growth followed a spatial pattern with significant growth in the vicinity of the *B. subtilis* colony (Figure 3a).

**Figure 3.**
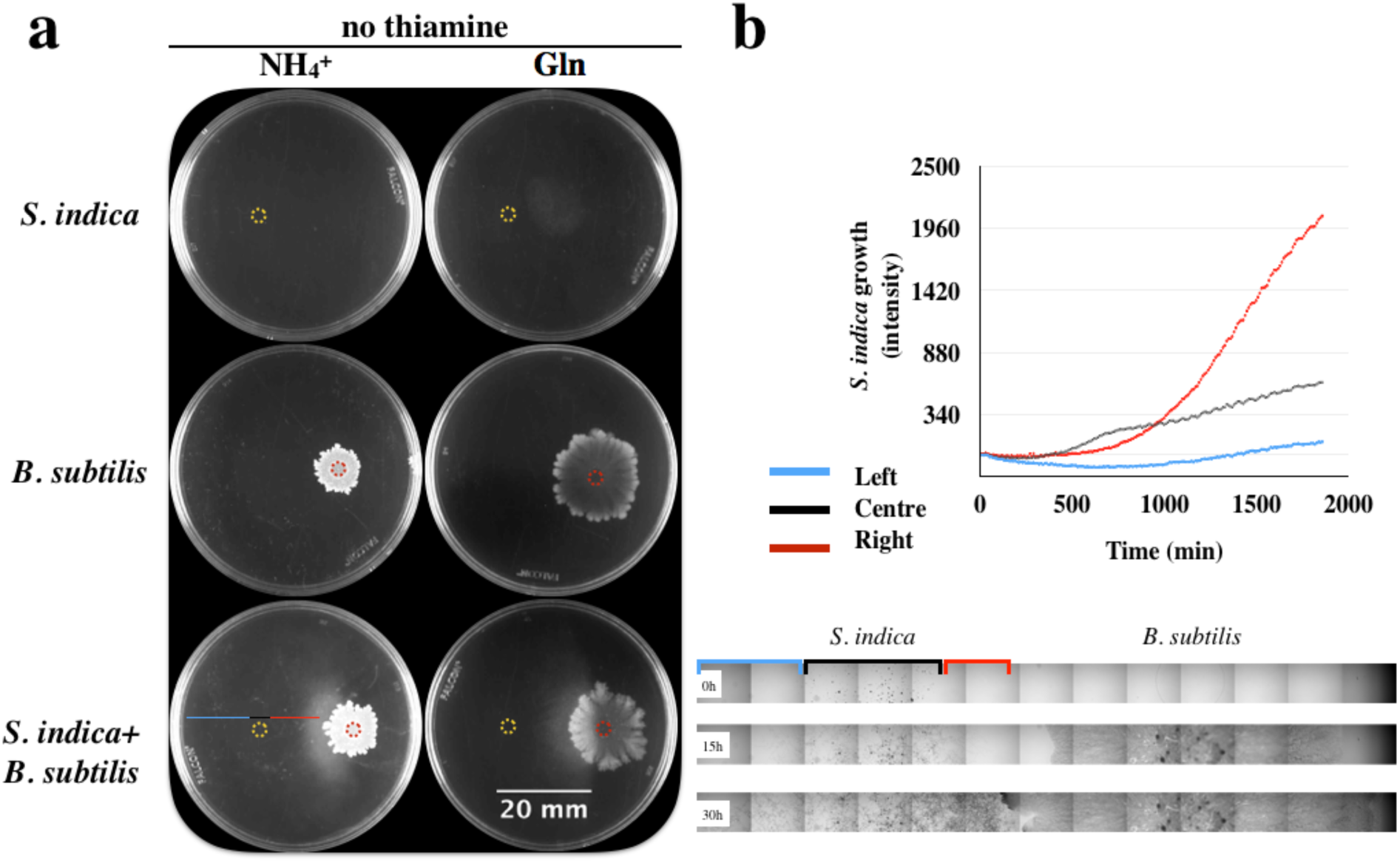
*S. indica* and *B. subtilis* interactions on agar plates. **a**. Rows from top to bottom showed growth of monocultures of *S. indica*, *B. subtilis*, and their co-culture respectively under the absence of thiamine. The yellow dotted circle on the images indicates the *S. indica* inoculation point. Red dotted circle indicates *B. subtilis* inoculation point. The left and right columns show growth on plates after two weeks, using ammonium and glutamine as nitrogen sources respectively. When both organisms were cultured together (bottom row), *B. subtilis* and *S. indica* were inoculated on the right and left of the plate respectively. Plates shown are representative of at least 3 replicates for each condition. We performed 2 biological replicates of this experiment, with qualitatively similar results. **b.** Bottom: Time lapse image series of *S. indica* and *B. subtilis* growth on agar plates in the absence of thiamine and with ammonium as nitrogen source. Each image strip is composed of microscopy images of the same horizontal section from the middle of the plate through the inoculation point of *S. indica* and *B. subtilis*, at 0 h, 15 h and 30 h after inoculation. *S. indica* growth across defined parts of this horizontal section (blue, black, and red lines – also compare bottom left plate in 3a) was quantified by measuring image density and plotted over time (upper chart). The sections correspond to the *S. indica* colony side closer to *B. subtilis* (red), the middle of the colony (black), and the colony side far from *B. subtilis* (blue).

We used time-lapse microscopy to quantify this spatial growth pattern of *S. indica* and found that growth (as approximated by image density) happened faster at the side closer to the *B. subtilis* colony compared to the far side of the plate (see Figure 3b and Supplementary Videos 3 and 4). This could be explained by the presence of an increasing thiamine gradient towards the *B. subtilis* colony that facilitates *S. indica* hyphal growth. While these findings strongly suggest a *B. subtilis*–linked thiamine provision, which then promotes *S. indica* growth, our attempts to quantify thiamine from agar plate co-cultures has failed, presumably due to a combination of thiamine consumption and sensitivity limitations of available thiamine assays (50 μg/L) (Lu and Frank, 2008). We were, however, able to detect thiamine from concentrated *B. subtilis* and found the concentration in liquid culture supernatants to be approximately 7.56 μg/l.

While the above findings strongly suggest that the growth enabling of *S. indica* by *B. subtilis* is due to thiamine supply, another theoretical possibility is that *B. subtilis* provides metabolites other than thiamine, that allow bypassing of central reactions requiring thiamine as a co-factor. In other words, provision of metabolites that are downstream of pyruvate in the TCA cycle (Figure 2b). To rule out this possibility, we have analyzed growth of *S. indica* in the absence of thiamine but supplemented with organic and amino acids that link to the central carbon metabolism. We found that none of the 17 amino acids or 8 organic acids tested or their combinations allowed for *S. indica* growth in the absence of thiamine (Figure S3). This finding further confirmed that *B. subtilis* facilitated growth of *S. indica* in thiamine-free medium is linked directly to thiamine.

### Metabolic profiling shows additional metabolic interactions between *S. indica* and *B. subtilis*

To analyse the basis of metabolic interactions between the two organisms and to collect more evidence for thiamine-based auxotrophy, we grew each organism in liquid culture on its own and then cross-cultured the other organism on the supernatant of the first one. As with agar plates, we found that in the absence of thiamine, the *S. indica* growth was limited to spore germination (Figure 4). When supplemented with *B. subtilis* supernatant, however, *S. indica* showed significantly increased growth in liquid culture (Figure 4). Consistent with this, there was also a growth enhancement of *S. indica* by the *B. subtilis* supernatant when cultured in the presence of thiamine.

**Figure 4.**
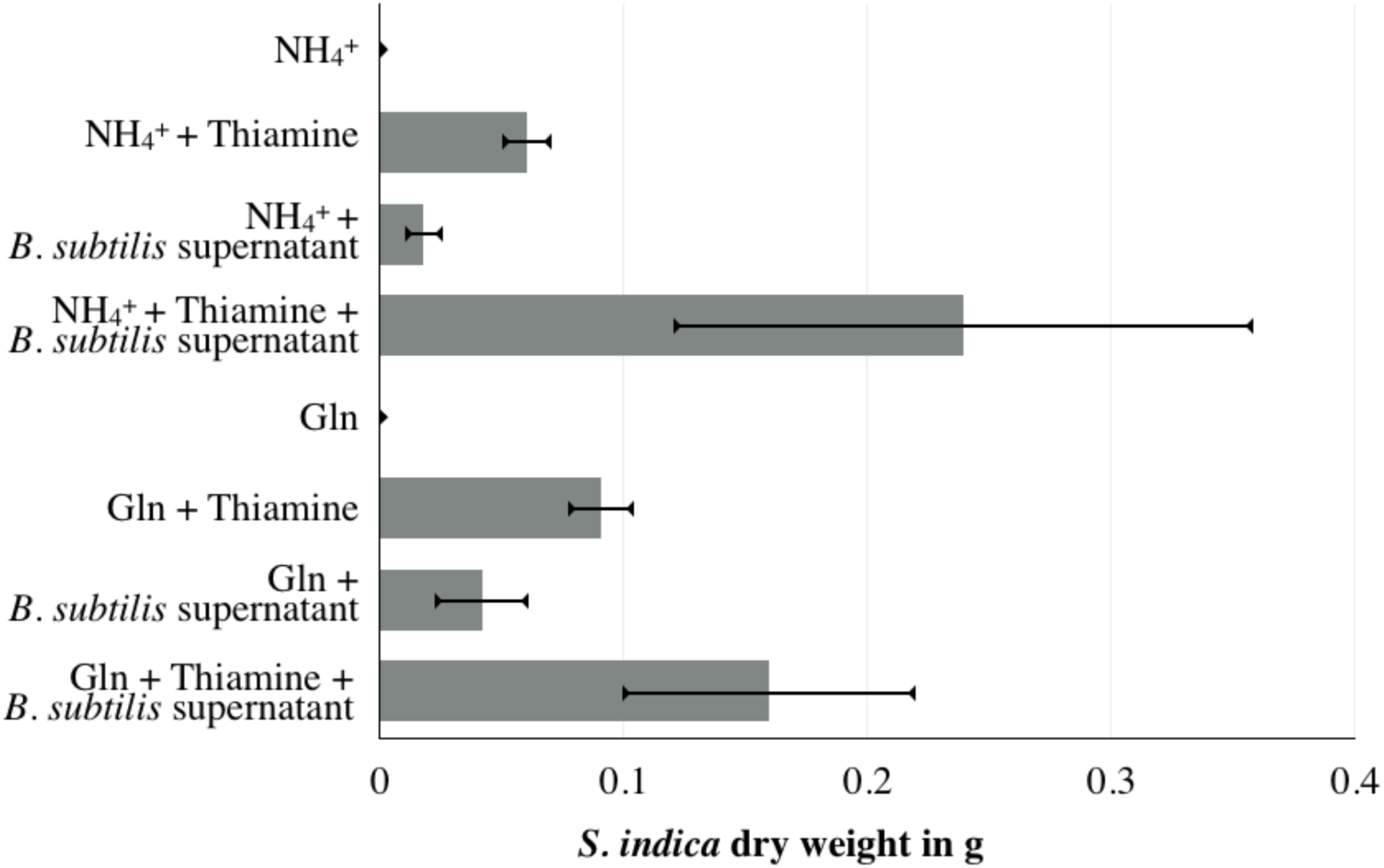
*S. indica* growth under different media compositions. Growth using either ammonia or glutamine and supplemented with thiamine or *B. subtilis* supernatant, as indicated on the y-axis. The x-axis shows *S. indica* growth approximated by total dry weight after one week of growth (see *Methods*).

To better understand the metabolic basis of these physiological observations, we repeated these experiments and quantified the concentrations of the key organic acids linking to the TCA cycle (lactate, acetate, pyruvate, and formate) in the supernatant of each organism before and after cross-cultivation using ion chromatography. We found that the supernatant from *B. subtilis* monoculture contained significantly higher amounts of acetate and some formate and pyruvate, and that the extracellular levels for these compounds did not change in the presence or absence of thiamine in the media (Figure S4). When *S. indica* was grown in the *B. subtilis* supernatant and in the absence of thiamine, it consumed both acetate and formate and produced pyruvate. In the presence of both thiamine and the *B. subtilis* supernatant, the consumption of acetate and formate was also observed, but there was also production of lactate in addition to pyruvate (Figure S4). These findings, in particular acetate and formate cross-feeding from *B. subtilis* to *S. indica*, explain the positive impact of *B. subtilis* supernatant on growth irrespective of thiamine availability. They also provide further support that the *B. subtilis-*associated growth of *S. indica* relates to thiamine provision rather than organic acids, since acetate and formate alone did not enable *S. indica* growth in thiamine-free media (Figure S3). We further found that *S. indica* secreted an unidentified organic acid in thiamine-media, that showed an IC profile overlapping with that of glutamine, and that was consumed by *B. subtilis* in the reverse experiment design (of growing *B. subtilis* in *S. indica* supernatant).

### The successful co-existence of *S. indica* and *B. subtilis* depends on spatiotemporal organization in the system

The above findings show that *B. subtilis* can support the growth of *S. indica* in thiamine-free medium either through its supernatant or when co-cultured at a distance on an agar plate. Both experimental setups were geared towards identifying possible interactions among the two species through utilization of the excretions of one species by the other, but did not necessarily consider the spatiotemporal factors on such interaction. Thus, a remaining question was whether both species could still co-exist and establish a successful interaction under different conditions regarding the spatial proximity or size of initial inoculation, or the actual growth phase that the different species are in at the time of introduction onto the agar. While addressing these questions is experimentally challenging, we attempted here to analyse the impact of spatial-temporal factors on the outcome of the *S. indica B. subtilis* auxotrophic interaction by changing inoculation time and location on agar plates. In particular, we separated the inoculation of *S. indica* spores from *B. subtilis* inoculation either in time (by inoculating spores 3 days before *B. subtilis* colony inoculation) or space (by inoculating *S. indica* and *B. subtilis* at certain distance to each other). Alternatively, we inoculated *S. indica* spores after mixing with *B. subtilis*. These experiments mimic a scenario commonly found in agri-technology practices when using pre-mixed cultures or spores of different microbes as soil biofertilizers or plant seed pre-treatments (Turner, 1991). We found that with temporal or spatial separation, both species could successfully grow in the absence of thiamine, indicating a positive auxotrophic interaction (Figure 5). In contrast, direct co-inoculation of *B. subtilis* with *S. indica* significantly hampered co-existence of the two species, particularly reducing *S. indica* growth (Figure 5). These findings indicate the significance of the spatiotemporal organization on microbial co-existence for the formation of stable interactions and, hence, on higher order microbial community structures. Our observations either reflect competition for resources upon co-inoculation, or alteration of the micro-environment by one species so that the other cannot establish itself, as discussed further below.

**Figure 5.**
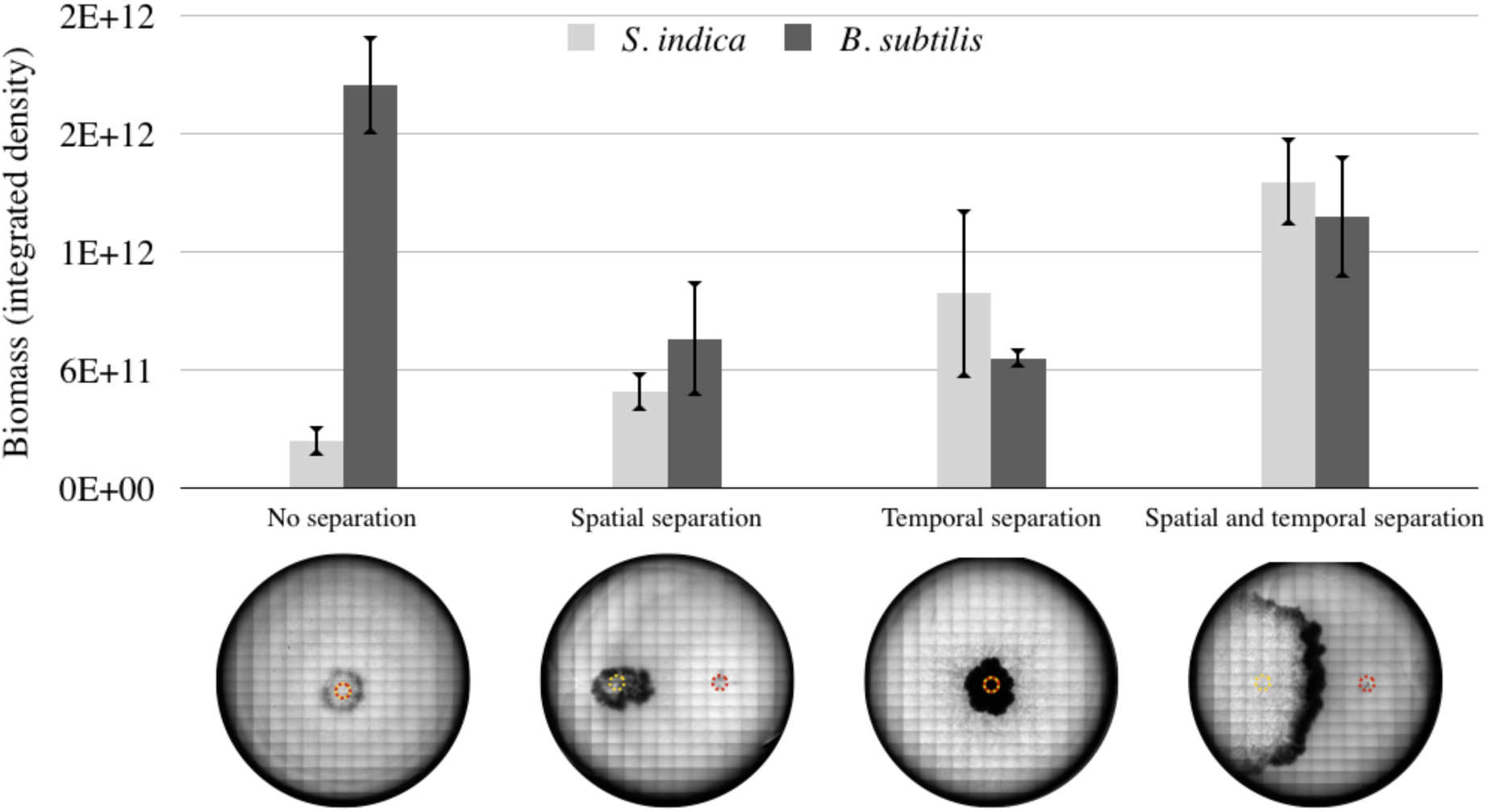
Biomass of *B. subtilis* and *S. indica* under different spatiotemporal culturing cases. “No separation” refers to *B. subtilis* culture and *S. indica spores* being pre-mixed at 1:1 volume ratio, and then inoculated as a single solution. “Spatial separation” refers to approximately 1.5 cm separation of *S. indica* (left) and *B. subtilis* (right) inoculation points. “Temporal separation” refers to inoculation of *S. indica* 3 days prior to *B. subtilis* inoculation. The yellow dotted circle on the images indicates the *S. indica* inoculation point. Red dotted circle indicates *B. subtilis* inoculation point. Growth of the different species was approximated by tracing their respective colonies on the plate and measuring the image intensity from the engulfing areas 2 weeks after *S. indica* inoculation. Measurements are from 3 replicate agar plates, with a representative plate image shown at the bottom. These images show microscopic scans of each plate at 2 weeks of growth. For the “no separation” case, there was no observable *S. indica* colony expansion after 1 week. We performed 2 biological repetitions of this experiment, with qualitatively similar results.

## Conclusion

Here, we report the discovery of a novel auxotrophy for thiamine in the endophytic fungus *S. indica* and how this thiamine requirement can be satisfied by *B. subtilis*. Our findings suggest that *S. indica* possesses thiamine transporters but lacks necessary thiamine biosynthesis genes, and its growth in thiamine-free medium cannot be supported by any other vitamin or relevant organic and amino acids. In contrast, co-culturing with *B. subtilis* in thiamine-free media allows *S. indica* growth, indicating a successful auxotrophic interaction between the two organisms. In addition, we have identified several cross-feeding interactions between the two organisms involving overflow and consumption of organic acids. We found that the auxotrophic interaction can only be achieved under conditions where the inoculation (and germination) of the two species is separated in time or space.

These findings have implications both for the study of *S. indica*, as an important plant-supporting soil fungus (Qiang *et al.*, 2012), and for the engineering and application of minimal synthetic communities that aim to establish plant supporting soil communities. In the former direction, future metabolic and physiological studies of *S. indica* will be enabled by the defined media conditions and identified thiamine auxotrophy in this study. A key suggestion from a biotechnological perspective, for example, is to consider thiamine as an important factor in the commercial mass production of *S. indica* (Singhal *et al.*, 2017). In the latter direction, the presented results point to the importance of the role of spatiotemporal dynamics on the outcome of microbial interactions. The failure of establishment of auxotrophic interactions between *S. indica* and *B. subtilis*, when co-cultured in the same place and time suggest that additional ecological factors can override naïve expectations from complementary metabolic interactions. This interplay between ecological forces, abundance, and interactions of auxotrophic species is currently not well-understood and only explored in a few studies, which used either synthetically engineered interactions (Shou *et al.*, 2007; Mee *et al.*, 2014) or enriched microbial communities (Embree *et al.*, 2015). Thus, our defined synthetic co-culture represents a powerful system towards understanding this new emerging axis of the molecular ecology of species interactions.

The observed failure to establish a successful auxotrophic interaction when co-cultured at the same time and space could be related to several factors. At the simplest level of explanation, *S. indica* and *B. subtilis* could be competing for and depleting the local carbon source. This is an unlikely explanation for our experiments, where carbon levels were relatively high. An alternative explanation would be that changes in ecological conditions caused by one species, would prevent the other species from establishing itself. In this ecological explanation, both the changes in local pH and in oxygen availability could be relevant. Changes in pH upon microbial growth were highlighted recently as affecting the subsequent microbial interactions and growth (Vylkova, 2017; Ratzke and Gore, 2017). Similarly, extensive oxygen depletion upon microbial colony growth on agar plates has been shown in several studies (Peters and Wimpenny, 1987; Dietrich *et al.*, 2013; Kempes and Okegbe, 2014) and can directly influence subsequent or simultaneous growth of different species or their interactions. This explanation could be particularly relevant in our experiments, where we expect *S. indica* germination and initial growth to require substantial oxygen (Tacon *et al.*, 1983).

The finding that the success of auxotrophic interactions relates to spatiotemporal effects suggest that consideration should be given to inoculation timing when designing or applying biofertilizers or bio-control agents to the soil. Indeed, microbial interactions and synergisms are suggested to be crucial for soil fertility, bioproductivity and ecosystem functioning (Perotto and Bonfante, 1997; Bulgarelli *et al.*, 2013; Pérez-Jaramillo *et al.*, 2016). Plants significantly benefit from symbioses with soil microbes, with benefits ranging from nutrient supply, growth promotion to elevating plant stress resistance (Vessey, 2003; Esser, 2013; Davison, 1988; Castillo *et al.*, 2013; Yurgel *et al.*, 2014). At the same time, soil microbes can interact among themselves or alter each other’s interactions with the plants (Veresoglou *et al.*, 2012; Lareen *et al.*, 2016) (Fitter and Garbaye, 1994; Kohlmeier *et al.*, 2005). The biochemical basis of these potential multi-level interactions in the soil has remained mostly elusive to date, with few documented cases of amino acid auxotrophies in specific soil bacteria and vitamin-provision from plants relating to their root colonization (Nagae *et al.*, 2016; diCenzo *et al.*, 2015; Streit *et al.*, 1996). The presented synthetic community of *S. indica* and *B. subtilis* shows for the first time that metabolic auxotrophy can directly underpin microbial interactions and growth, and that the success of interaction can be determined by spatiotemporal organization in the system. In to the future, this synthetic system allows controlled investigation (and potential optimization) of fungal-bacteria interactions and can be further extended with additional microbes and a plant. The resulting minimalist synthetic eco-system can provide a platform to analyze and control cross-kingdom relationships between plants and their growth promoting fungi and bacteria, and enable new applications for vertical farming and crop production in the field.

## Materials and methods

### *S. indica* cultures, growth media and conditions

The defined basic medium for testing *S. indica* growth with ammonium as nitrogen source contained per liter; 15g agar, 20g glucose, 1.32 g (NH_4_)_2_SO_4_, 0.89 g Na_2_HPO_4_·2H_2_O, 0.68 g KH_2_PO_4_, 35 mg Na_2_MoO_7_·2H_2_O, 5.2 mg MgSO_4_, 2.5 mg FeSO_4_·7H_2_O, 0.74 mg CaCl_2_·2H_2_O, 0.0043 mg ZnSO_4_·7H_2_O, 0. 004 mg CuSO_4_·5H_2_O. Growth experiments for testing effects of different vitamins were performed in 24-well plates (Ref: 353226, Falcon), where each well contained 2 ml of the basic medium, supplemented with either 200 μg/l of single vitamin solution, 1 g/l yeast extract, 200 μg/l mixture of all eight vitamins tested, or equivalent amount of water. Each well was then inoculated with 1 μl of *S. indica* spore suspension (approximately 500,000 spores/ml, counted with Neubauer counting chamber), where spores were harvested from 6-8 weeks old *S. indica* agar plates. In the ‘non-inoculated’ control treatment, 1 μl of water was used instead. Each treatment condition was prepared in three technical replicates. 24-well plates were then sealed with parafilm and placed in a 30 °C static incubator for 2 weeks. Images were taken with a gel doc system (Syngene) at the end of this period.

### Experiments on agar plates, growth media and conditions

The defined (basic) medium for testing *S. indica* growth on agar plates contained per liter; 15 g agar, 20 g glucose, 0.89 g Na_2_HPO_4_·2H_2_O, 0.68 g KH_2_PO_4_, 35 mg Na_2_MoO_7_·2H_2_O, 5.2 mg MgSO_4_, 2.5 mg FeSO_4_·7H_2_O, 0.74 mg CaCl_2_·2H_2_O, 0.0043 mg ZnSO_4_·7H_2_O, 0.004 mg CuSO_4_·5H_2_O. When the chosen nitrogen source was ammonium, 1.32 g/l (NH_4_)_2_SO_4_ was added to this basic recipe. When the chosen nitrogen source was glutamine, 1.46 g/l glutamine was added to this basic recipe. For experiments with thiamine, 150 μg/l thiamine was added to the basic media after autoclaving.

Experiments were carried out on 60 mm dishes, filled with 6ml of agar medium given above. A 500,000 spores/ml *S. indica* spore suspension, with spores harvested from 6-8 weeks old *S. indica* agar plates, was inoculated with 2 μl (on the left side of the plates). At approximately 2 cm distance to the right of the inoculum, either a ‘mock’ solution, a thiamine block, or a *B. subtilis* inoculum were placed. The ‘mock’ solution was 2 μl of sterile water. The thiamine blocks were made by pouring 6 ml of 1.5% agar solution containing150μg/l thiamine into a 60 mm plate, and then punching a block out using a sterile pipette tip with a diameter of 4.7 mm. Therefore, each agar block used contained approximately 5.6 ng thiamine. The *B. subtilis* inoculum was a 2 μl sample harvested from a culture, grown in 5ml liquid Lysogeny broth (LB) (Bertani, 1951) to an OD_600_≈ 0.5 measured by spectrophotometer (Spectronic 200, Thermo Fisher Scientific), then washed and re-suspend to OD_600_ 0.5 with 10 mM MgCl_2_. The plates were incubated in 30 °C for 2 weeks. Images were taken with gel doc system (Syngene) at the end of this period.

### Time-lapse microscopy and image analysis

Time-lapse microscopy was performed on agar-medium cultures that were prepared using the same basic medium described above. A 6-well tissue culture plate (Ref: 353046, Falcon) was used and 1 μl of *S. indica*, *B. subtilis* or mock solution prepared as described above were inoculated on each well accordingly to experiment design. An Olympus IX83 microscope, UPlanFLN 4× objective and cellSens software were used for recording the growth. Okolab stage top incubator (H301-T-UNIT-BL-Plus system, and H301-EC chamber) were used for incubation, with a temperature sensor and lens heater set to 30 °C and stabilized for at least 2 hours prior to the experiment. Different fields of view were chosen at interior and periphery of each colony and images from those fields were recorded using the automated microscope stage and Olympus cellSens software. Images were taken in 1 hour intervals and put together as image series. ImageJ (Fiji) (Schindelin *et al.*, 2012) was used for measuring the mean intensity on each field of view over time, normalized against the intensity value of the first frame from each view point (as shown in Figure 3).

### Spatial and temporal separation experiments

*S. indica* (500,000 spores/ml, determined by counting with Neubauer counting chamber) and *B. subtilis* (OD_600_≈ 0.5, determined by spectrophotometer (Spectronic 200, Thermo Fisher Scientific)) were cultivated on thiamine-free synthetic medium containing ammonium as sole nitrogen source. 6-well tissue culture plates (Ref: 353046, Falcon). On each plate, 1 μl of *S. indica* and 1μl of *B. subtilis* were inoculated on 5 of the wells; one well was intentionally left non-inoculated as a blank. In the “no separation” case, *S. indica* and *B. subtilis* were pre-mixed at 1:1 volume ration, and inoculated on the center of each well. For “spatial separation” case, *S. indica* was inoculated 7.5 mm left to the center of a well and *B. subtilis* 7.5mm right to the center, leaving 15 mm distance in between. For “temporal separation” case, *S. indica* was inoculated on each well, the plates were then incubated for 3 days and *B. subtilis* was inoculated after this time. All the plates were incubated in 30°C for 2 weeks (starting from the time of *S. indica* inoculation). Images were taken by scanning each well under a microscope (Olympus IX83) using the same exposure time under bright field.

ImageJ was used for measuring the biomass by integration of the total colony density. An image of each colony was manually outlined using the selection tool. The selected area was compared with the same location on a blank well from the same plate. The area and relative intensity were recorded (using “measure” function) and used for calculating the colony growth.

### Supernatant cross-feeding experiments

Axenic cultures of *S. indica* and *B. subtilis* were cultivated in 50 ml basic medium described above, with or without thiamine. For *S. indica* an inoculum of 50 μl of a 500,000 spores/ml spore suspension, harvested from 6-8 weeks old *S. indica* agar plates, was used. For *B. subtilis*, an inoculum of 50μl, sampled from a culture grown in LB to an OD_600_≈ 0.5 determined by spectrophotometer (Spectronic 200, Thermo Fisher Scientific), and washed with 10 mM MgCl_2_, was used. After one week incubation in 30°C and at 150 rpm, *S. indica* cells were harvested by centrifugation at 18,000 g for 20 minutes. The supernatant was collected and filtered through a 0.2 μm polyethersulfone (PES) filter (Ref: WHA67802502, Whatman), while biomass was washed with 40 ml MilliQ water, dried using a centrifugal evaporator (EZ-2 Elite, Genevac), and then weighted. The growth of *B. subtilis* liquid cultures was monitored daily by taking 1 ml samples and measuring OD_600_ by spectrophotometer (Spectronic 200, Thermo Fisher Scientific). At the end of 1 week, the remaining liquid culture was centrifuged at 18,000 g for 10 min. The supernatant was collected and filtered through a 0.2 μm PES filter, while biomass was discarded.

Both supernatants were mixed with fresh basic medium in a 1:1 ratio to set up new axenic cultures of *S. indica* and *B. subtilis*. 1 ml liquid samples were collected from these cultures after one week by filtering through a 0.2 μm nylon membrane (Ref: WHA7402004, Whatman). Samples were transferred into polypropylene vials (Ref: 079812, Thermo Fisher Scientific) for Ion Chromatography, which was performed using Dionex ICS-500+ and column Dionex IonPac AS11-HC-4 μm (2 x 250 mm).

### Sequence analysis and BLAST search

Sequences of key thiamine biosynthesis enzyme genes (Table S1) from *Saccharomyces cerevisiae* S288c were compared with available *S. indica* genome homologues using Position-Specific Initiated BLAST (https://blast.ncbi.nlm.nih.gov/Blast.cgi), to identify the putative functions of the corresponding genes. An e-value of 1e-6 was chosen as cut-off to identify homologous sequences (Altschul, 1997). The results of the analysis using blastx are shown in Table S1, while blastn and tblastx did not return any results with the chosen cut-off.

### Thiamine measurements on *B. subtilis* liquid cultures

Axenic cultures of *B. subtilis* were cultivated in 50ml synthetic medium containing glutamine as sole nitrogen source without thiamine. *B. subtilis* inoculum of 50 μl, sampled from a culture grown in LB to OD_600_≈ 0.5 determined by spectrophotometer (Spectronic 200, Thermo Fisher Scientific), and washed with 10 mM MgCl_2_ was used. After one week of incubation in 30 °C and at 150 rpm, *B. subtilis* cultures were harvested by centrifugation at 18,000 g for 10 minutes. The supernatant was collected and filtered through a 0.2 nm PES membrane filter (Ref: WHA67802502, Whatman). The supernatant of 250 μl from each culture was transferred to a clean 1.5 ml Eppendorf tube, followed by sequentially adding 10 μl 1% K_3_[Fe(CN)_6_], 150μl 15% NaOH solution and 150μl isobutanol. The tubes were shaken vigorously for 1 minute, followed by 2 minutes of centrifugation at 13,000 g. The upper isobutanol layer of each tube was transferred to a new 1.5 ml Eppendorf tube, containing approximate 0.2g Na_2_SO_4_. The tubes were mixed well and centrifuged for 1 minute at 13,000 g for solids to settle. 100 μl of supernatant from each tube were transferred to 96-well plates (Ref: 3916, black flat bottom, Corning) and the fluorescence was measured using a plate reader (CLARIOstar, BMG Labtech) at 365 nm excitation and 450 emission. The concentration of thiamine was determined with a series of known concentration standard thiamine solutions under the same treatment.

### *S. indica* growth under different thiamine concentrations

Synthetic medium containing ammonium as sole nitrogen source was used for testing *S. indica* growth in different thiamine concentrations. Media containing final thiamine concentrations of 0 μg/l, 1.5 μg/l, 15 μg/l, 150 μg/l and 1500 μg/l were prepared, and distributed in a 6-well tissue culture plate (Ref: 353046, Falcon). Each 6-well plate contained one concentration condition, with 3ml medium in each well. On each plate, 1μl of *S. indica* (500,000 spores/ml) were inoculated on the center of five wells, while one well was intentionally left non-inoculated as a blank. Plates were incubated at 30 °C for 2 weeks. Afterwards, lids were removed and OD_600_ and fluorescence at 365 nm for excitation and 450 nm for emission were measured using a plate reader (CLARIOstar, BMG Labtech) and the plate scan function to get an overall reading of each well. Images of plates were taken using a gel doc system (Syngene).

## SUPPLEMENTARY FIGURES

**Supplementary Figure 1.**
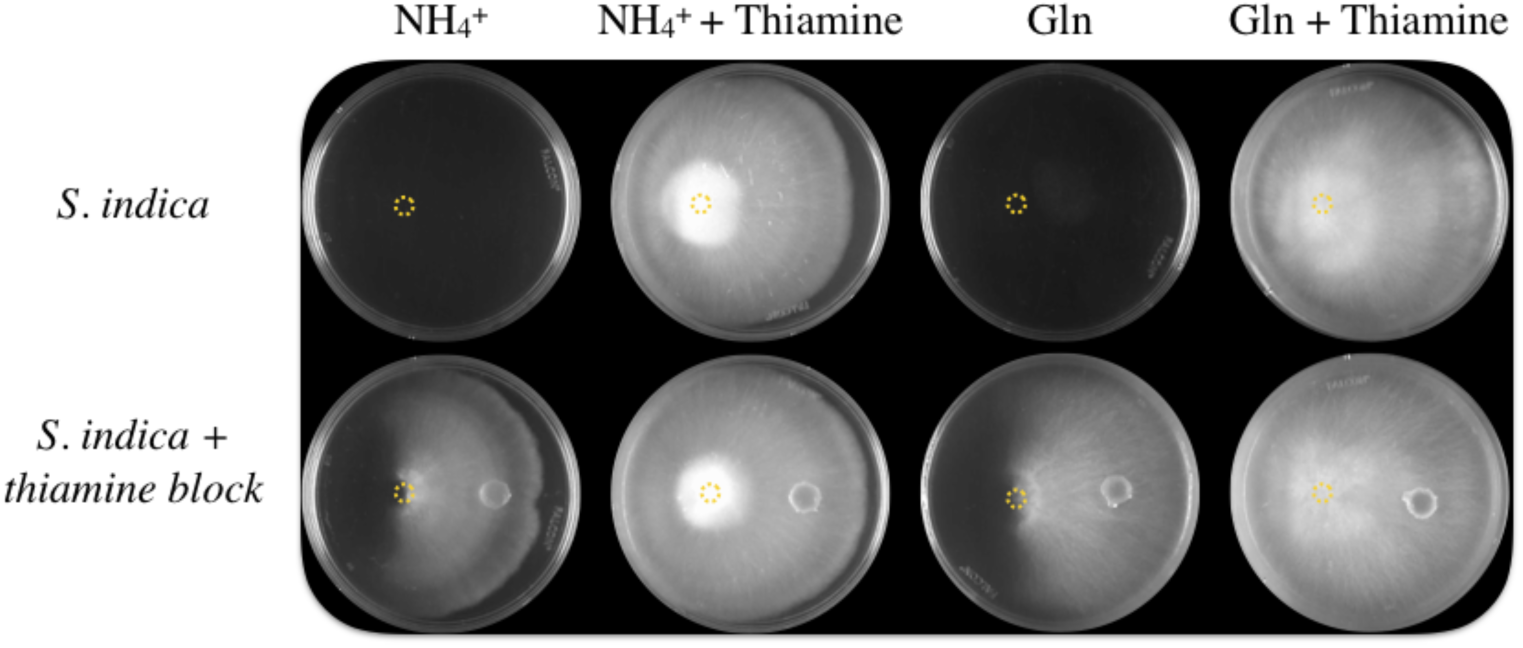
*S. indica* growth on agar plates after two weeks. Columns show different media containing thiamine or not and using ammonium and glutamine as nitrogen sources (shown as “NH4+” or “Gln” on the panels). The top and bottom rows show the images of agar plates without or with an additional agar block containing thiamine, placed ∼1.5 cm to the right side of *S. indica* inoculation point, which is indicated with a yellow circle.

**Supplementary Figure 2.**
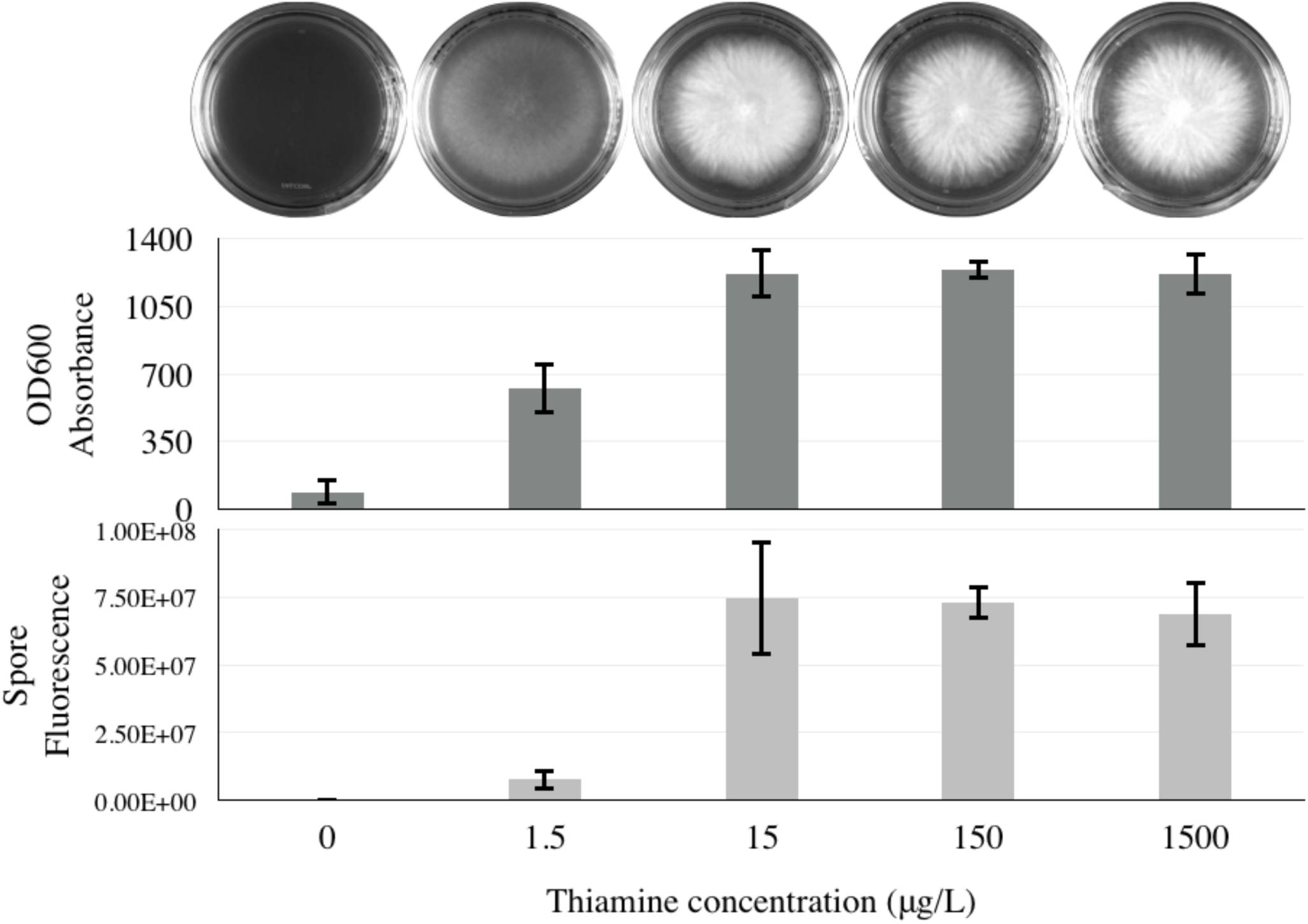
*S. indica* growth on medium containing different concentrations of thiamine and ammonium as nitrogen source. Images at top show two weeks growth of *S. indica* on agar plates, and at different concentrations of thiamine as shown below on the bottom *x*-axis. Each condition was repeated 6 times and images shown here are representatives for each condition. Upper and lower bar-plots show plate absorbance at OD_600_ and fluorescence intensity (measured at 390 excitation and 470 emission for detection of *S. indica* spores) respectively.

**Supplementary Figure 3.**
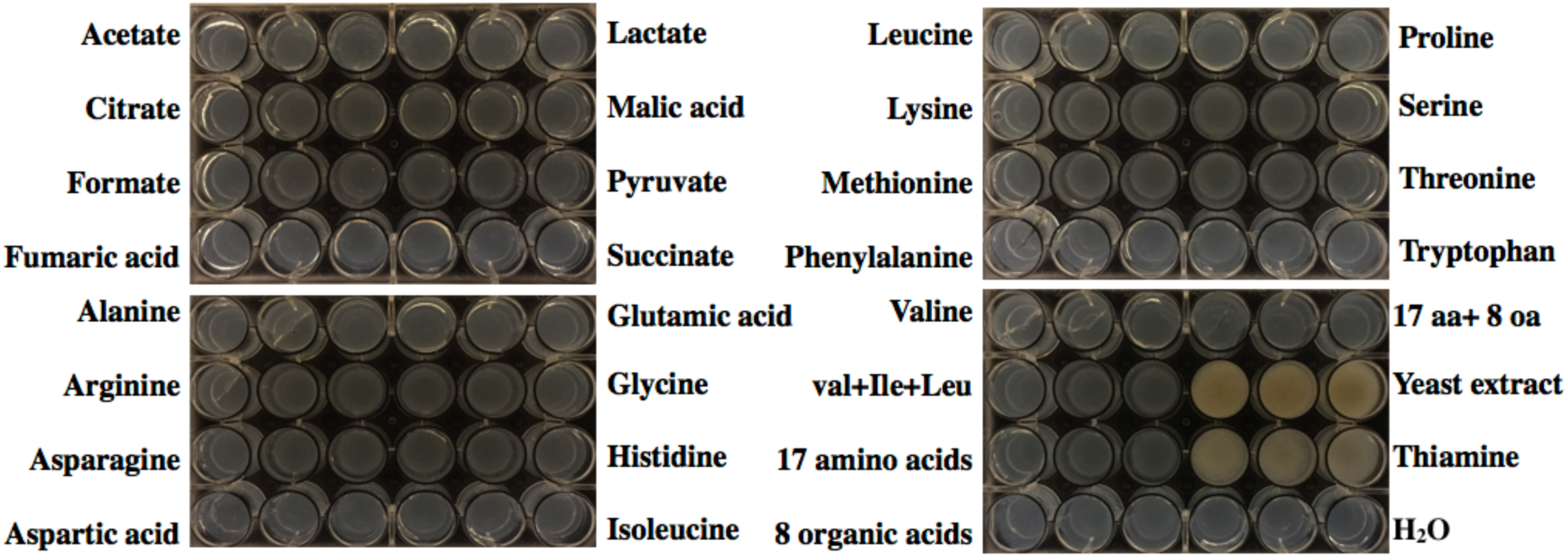
*S. indica* amino acid and organic acid screen. *S. indica* growth on the media containing different amino acid or organic acid as supplement. Each treatment has three replicates presented in 3 adjacent wells. *S. indica* growth is visible as white-yellow colonies on the surface of a well. Images were taken after two weeks of growth.

**Supplementary Figure 4.**
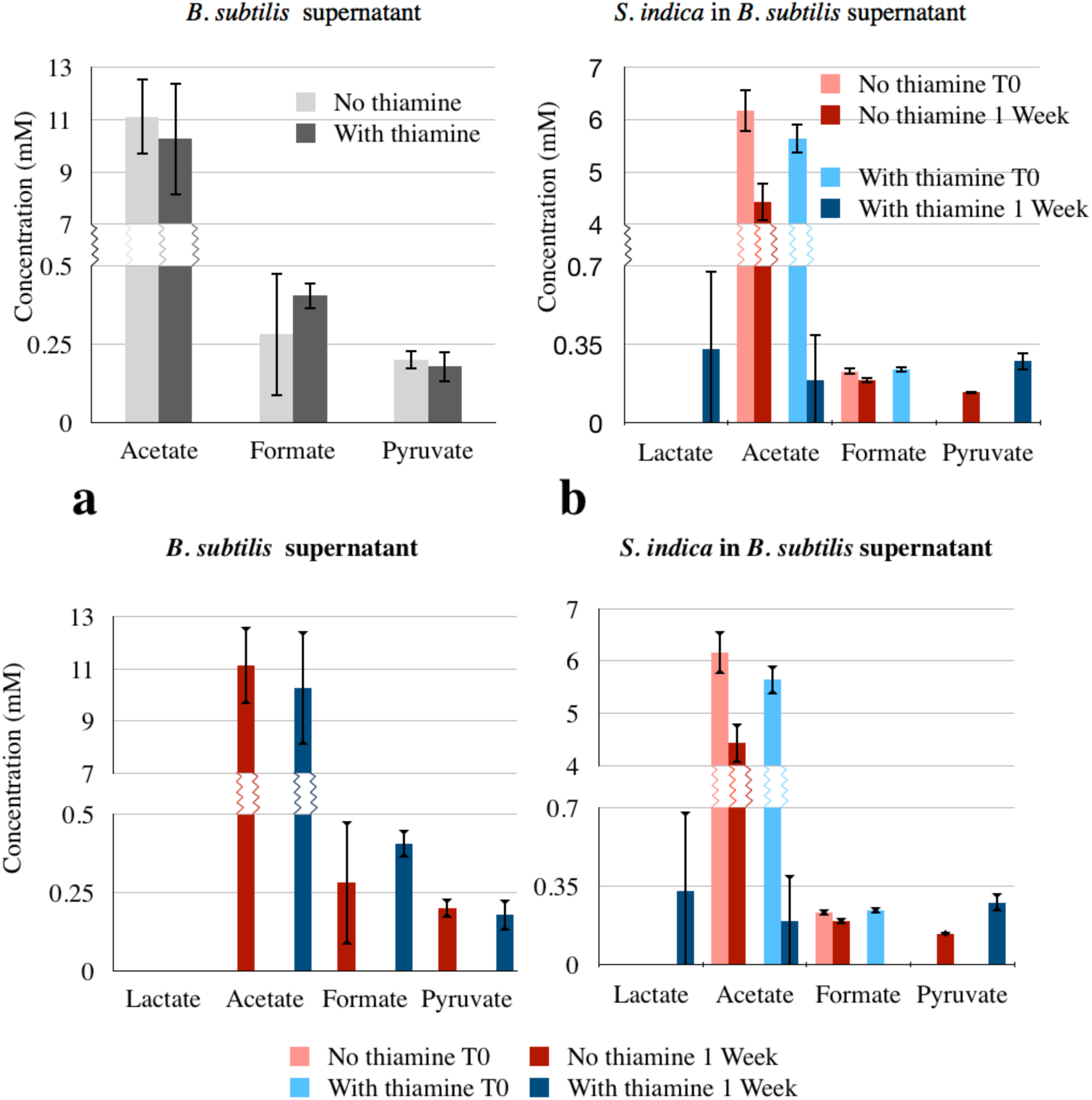
Concentrations of key organic acids in the supernatant of *B. subtilis* growth media in the presence of absence of *S. indica*. The left chart shows the supernatant after one week of *B. subtilis* cultivation in liquid medium containing glutamine as sole nitrogen source. The right chart shows the supernatant after one week of cultivation of *S. indica* in *B. subtilis* supernatant mixed with the same volume of fresh medium. T0 and T1 indicate the initial condition and after 1 week of growth respectively. Note that T0 concentrations in the right chart correspond to concentrations in the left chart diluted with fresh media.

**Supplementary Table 1.** <u>Comparison of the thiamine-related genes/proteins of</u> *<u>S. indica</u>* <u>wit</u>h those of *S. cerevisiae*

| Gene name | Function | Source organism for sequence | Blastx result using *S.indica* genome<br>Max score/Query covery/E value |
| --- | --- | --- | --- |
| THI6 | Bifunctional hydroxyethylthiazole kinase/thiamine-phosphate diphosphorylase | *Saccharomyces cerevisiae* S288c | No |
| THI80 | Thiamine diphosphokinase |  | 104/84%/4e<sup>-26</sup> |
| Thi4 | 1. Thiamine thiazole synthase<br>2. Mitochondrial DNA damage tolerance |  | 47.8/15%/2e<sup>-06</sup> |
| THI5 | Pyrimidine precursor biosynthesis |  | No |
| THI11 | Pyrimidine precursor biosynthesis |  | No |
| THI12 | Pyrimidine precursor biosynthesis |  | No |
| THI13 | Pyrimidine precursor biosynthesis |  | No |
| PHO3 | Thiamin-repressible acid phosphatase |  | 114/86%/3e<sup>-27</sup> |
| THI10 (THI7) | Thiamin transporter |  | 176/92%/5e<sup>-48</sup> |
| PDC2 | Pyruvate decarboxylase, transcriptional activator in response to thiamin starvation. |  | No |
| RPI1 | Transcriptional regulator |  | No |

**Supplementary video 1.** *S. indica* monoculture on synthetic medium without thiamine.

**Supplementary video 2.** *S. indica* monoculture on synthetic medium with thiamine.

**Supplementary video 3.** *S. indica* and *B. subtilis* co-culture on synthetic medium without thiamine.

**Supplementary video 4.** *S. indica* and *B. subtilis* co-culture on synthetic medium with thiamine.

